# Cross-order detection of bacteriophage transduction in communities using ribosomal RNA barcoding

**DOI:** 10.1101/2025.05.03.652062

**Authors:** Zachary W. LaTurner, Matt J. Dysart, Samuel K. Schwartz, Elizabeth Zeng, James Chappell, Jonathan J. Silberg, Lauren B. Stadler

## Abstract

Bacteriophages (phages) facilitate gene transfer and microbial evolution in all ecosystems and have applications as tools for engineering microbiomes and as antimicrobials. Historic efforts to map phage hosts, such as plaque assays, are limited to culturable bacteria, are low throughput, and are hard to apply in environmentally-relevant contexts. To overcome these limitations, a synthetic ribozyme that stores information about DNA uptake in 16S ribosomal RNA (rRNA) was used to identify phage-host interactions by integrating the ribozyme into phage-plasmid P1 and performing targeted 16S rRNA sequencing following transduction. Experiments in synthetic and wastewater communities revealed Aeromonadales as a novel P1 host order and transduction of P1 into pathogens. Host range varied across phagemids with different origins of replication and phage particles with different tail fibers. This work shows how autonomous barcoding can be used in phages to identify the molecular controls on their host range in communities.

## Main

Bacteriophages (phages) play a crucial role in the environment. They drive ecological interactions in microbial communities by killing their hosts, conferring fitness costs on survivors, and supplementing the suite of functional genes at their hosts’ disposal.^1,2^ At an estimated abundance of 10^31^ viral particles on Earth, viruses are the most abundant lifeform and an order of magnitude more abundant than bacteria.^3^ Phages can act as agents of horizontal gene transfer (HGT), functioning as both disseminators of antibiotic resistance genes and as vectors for *in situ* gene delivery.^4–8^ Phages that kill pathogens also represent a promising alternative to antibiotics, as antibiotic-resistant infections are projected to kill up to 10 million people per year by 2050.^9^ Connecting phage to their different microbial hosts is essential for harnessing phages for therapeutic applications and understanding their ecology. Conservative diversity estimates suggest 10^7^-10^9^ distinct viral species, and given that many viruses infect more than one host, this suggests an even greater number of unique virus-host interactions.^10^

Experimental approaches for mapping phage-host interactions are limited in their ability to study transduction in communities. For example, plaque assays can only be applied to plaque-forming phages and culturable microbes.^11^ Viral tagging and electrophoretic methods overcome this challenge by isolating phage bound to bacteria or bacterial cell wall components, but cannot differentiate between phage absorbed to bacterial surfaces and phage that infect cells.^12,13^ Visualization methods that use fluorescence in situ hybridization to observe transduction events are hampered by complex methodology and the low efficiency of fluorescent probe delivery to cells.^14^ Approaches that link genetic material co-located in a cell, like epicPCR and high-throughput chromosome conformation capture (Hi-C), require intensive and sensitive sample processing to link phage and host DNA.^15–17^ Environmental transformation sequencing (ET-seq) avoids this processing by autonomously barcoding host genomes.^18^ However, ET-seq only yields signals in cells that transcribe and translate the synthetic components that modify chromosomes, and requires metagenomic sequencing, limiting throughput and affordability. Thus, there is a need for simpler experimental approaches that overcome these limitations.

Recently, a synthetic biology tool for studying HGT was reported that allows for sensitive detection of host range.^19^ This approach, called RNA-addressable modification (RAM), uses an engineered ribozyme to write information about participation in conjugation to 16S rRNA of recipient cells. The ribozyme is transcribed using a single broad host range promoter and consists of a guide sequence targeting 16S rRNA, a catalytic core, and a synthetic barcode. The guide sequence directs the ribozyme to a conserved region of 16S rRNA, where it catalyzes a trans-splicing reaction that adds the barcode adjacent to variable regions in 16S rRNA that can be used for taxonomic identification (Fig. 1a). When used to monitor plasmid host range via conjugation in wastewater, RAM revealed plasmid uptake by ∼50% of the amplicon sequence variants (ASVs).^19^ When coded into multiple plasmids, RAM revealed how changes in replication origins altered conjugative host range. These observations suggest that RAM could be used with other types of mobile DNA, like phage, to simplify host range studies.

**Figure 1.**
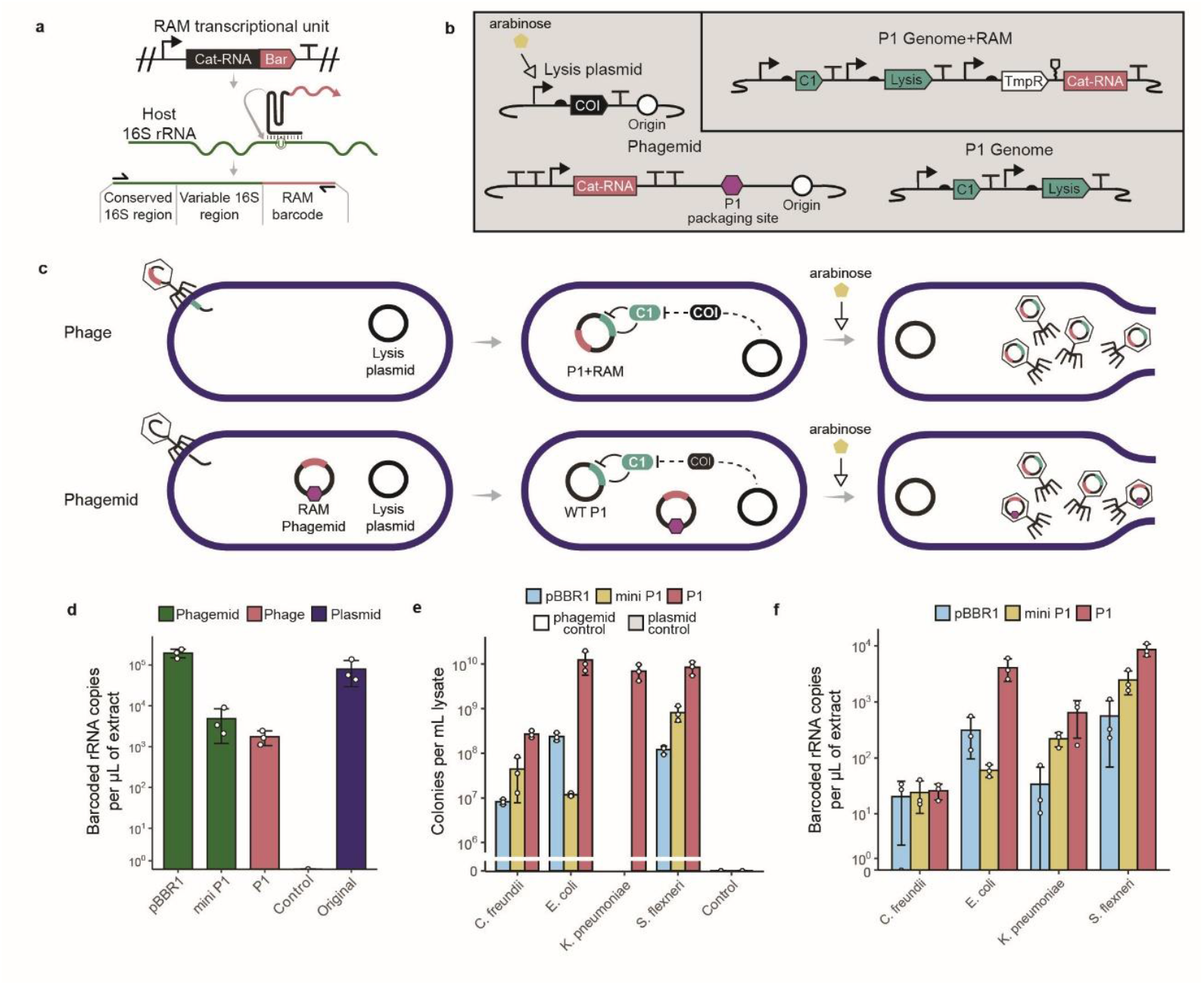
Benchmarking RAM in the phage P1. **a**, Mechanism of RAM barcoding. Transcription of the RAM operon produces a ribozyme that catalyzes the splicing of a barcode onto host 16S rRNA. Primers for the spliced product can capture the total amount of barcoded 16S rRNA through RT-qPCR and/or allow for sequencing of the conserved and variable regions of 16S rRNA adjacent to the barcode. **b**, Genetic elements deployed in the phage and phagemid experiments. Genomically integrated RAM (P1 genome + RAM) includes TmpR in the RAM operon to allow for selection and maintenance of modified lysogens. Phagemids include a P1 packaging site so they will be incorporated into phage particles. The lysis plasmid contains the c one inactivator (*coi*) gene under arabinose inducible expression. Coi represses the C1 repressor, which itself represses the lytic cycle of P1. Addition of arabinose induces the lytic cycle of P1 in this system. **c**, Process for the production of phage particles containing RAM constructs. Top: *E. coli* MG1655 containing the lysis plasmid is lysogenized with P1 phage containing RAM in the genome. Addition of arabinose produces phage particles, all containing P1 genome with RAM. Bottom: *E. coli* MG1655 containing the lysis plasmid and the phagemid is lysogenized with P1 phage. Addition of arabinose produces a mixture of phage particles, some containing phagemid and some containing P1 genome without RAM. **d**, Quantification by RT-qPCR of barcoded 16S rRNA in *E. coli* MG1655 when all cells in the population contain a given phagemid (pBBR1, miniP1), phage with RAM (P1), phage without RAM (Control), or original pBBR1 plasmid from a prior study (Original).^19^ **e**, CFUs generated through selective plating of different species transduced with phage particles containing the given construct. Phage control involved infection of *E. coli* MG1655 with wildtype P1 phage particles lacking the RAM operon. Phagemid control involved infection of *E. coli* MG1655 by lysate generated from a host without a phagemid but a modified version of P1 with KanR in the genome. **f**, Quantification by RT-qPCR of barcoded 16S rRNA produced through transduction of different species with phage particles containing the given construct. For panels d, e, and f, All data was produced with three biological replicates. Points indicate individual replicates, bars indicate the average value, and error bars the standard deviation.

To connect phages with hosts in a microbial community, we coded RAM into P1 phage and assessed transduction by sequencing barcoded-rRNA. P1 was targeted because it is a lysogenic phage that exists as an extrachromosomal element during the lysogenic phase of its life cycle; these phage-plasmids cannot be connected to hosts via metagenomic sequencing.^20,21^ P1 host range is important to understand because it infects many human-associated enteric bacteria and similar phages are thought to contribute to the spread of antibiotic resistant infections.^22–26^ By coding RAM in phage P1 and phagemids, we demonstrate RAM as a viable technology for reporting on phage-host relationships in diverse contexts. We also discover novel P1 hosts when performing studies in synthetic and wastewater communities. Finally, we investigate how two different tail fibers natively encoded in the P1 genome contribute to host range. This study represents a blueprint for uncovering the determinants of phage host range, and engineering phages for high efficiency DNA delivery for microbiome engineering.

## Results

### Phage genomes and phagemids engineered with RAM connect phages to their host

To deploy RAM in phages and record information on phage-host connections, we first added RAM to phage P1 in multiple genetic contexts (Fig. 1b, 1c). Using a CRISPR-associated transposon system (CAST), we inserted RAM into the genome of P1 (Fig. S1a).^27,28^ We also incorporated RAM into P1 phagemids, which are synthetic plasmids that contain the P1 phage packaging site (Fig. S1b).^25,29^ We created two phagemids with RAM, one containing the broad host range origin of replication (oriR) pBBR1, and one containing the native P1 oriR (referred to here as “mini P1”). pBBR1 is a broad host oriR and thus could be used for gene delivery where the goal is to stably maintain phagemid DNA in diverse hosts. Additionally, we envisioned the mini P1 phagemid would have a host range close to that of the full P1 phage but with the benefit of being easier to manipulate than the P1 genome, enabling additional studies on P1 host range. Phagemids are also unable to mobilize themselves without the help of other genetic elements, a benefit in the context of biocontainment.

We next delivered the RAM-containing phage and phagemids to *Escherichia coli* MG1655, via transformation for the phagemids and transduction for the phage, and used a selectable marker to obtain a population of cells that all contained the given construct. We observed barcoded 16S rRNA with all constructs, (Fig. 1d), demonstrating that RAM encoded on both the phage genome and phagemids generates a quantifiable RAM signal. We found that incorporating a packaging site into our previous, pBBR1 based plasmid to create a phagemid slightly affected RAM signal (Welch’s, two-tailed, unpaired t-test. *P* = 0.043 pBBR1 != Original). The P1 based constructs led to a decrease in signal relative to the pBBR1 phagemid, which we propose is due to a lower plasmid copy number of these constructs in *E. coli* (Welch’s, one-tailed, unpaired t-test. *P* = 0.01, 0.01 for mini P1 < pBBR1 and P1 < pBBR1, respectively).

To assess whether P1 phage and phagemids with RAM function in other strains, we generated phage particles containing the individual constructs and introduced them to different enteric bacteria previously demonstrated to be susceptible to P1 infection.^25^ In addition to *E. coli*, we tested *Citrobacter freundii, Klebsiella pneumoniae*, and *Shigella flexneri*. Unlike the previous experiment, this experiment included an infection step leading to a heterogenous population of cells with and without DNA constructs. We quantified infection by selective plating using antibiotic resistance genes on the phagemid or phage genome as well as by measuring RAM output via RT-qPCR (Fig. 1e, 1f). Preliminary experiments showed no dilution of phage particles (apart from the 1:10 dilution caused by addition to the bacterial culture) and a two hour incubation time optimized RT-qPCR signal with our protocol. (Fig. S3, S4)

We found that the genomically integrated RAM (P1) consistently produced the most CFUs across all species determined via selective plating (Welch’s, one-tailed, unpaired t-test. P < 0.05 for all P1 > construct comparisons. See Fig. 1e data). This is likely because the lysate of phagemid constructs (pBBR1 and mini P1) consists of a mixture of phagemid containing viral particles and wildtype P1-genome containing particles. Thus, only a fraction of the total phage particles contain RAM in the phagemid context, whereas in the genomically integrated context, all phage particles contain RAM (Fig. S2b). The phagemid production process also reduces the total amount of phage particles produced regardless of whether they contain a phagemid or wildtype P1-genome (Fig. S2a). The relative rank of phagemid CFUs within a species was not consistent across species. This may be because the different species have different defense systems with different degrees of efficacy against genetic sequences or proteins contained in the constructs. CFU counts and RT-qPCR signal generally agreed, despite RT-qPCR based measurements only requiring transcription coupled with a splicing reaction, while selective plating requires transcription and translation. This provides an important validation of RAM against widely used methods for quantifying phage transduction. Additionally, we could not quantify CFUs for phagemids in *K. pneumoniae* because it was already resistant to the selectable marker (AmpR), illustrating a benefit of using RAM to quantify infection over selective plating-based methods, particularly in microbes with native resistances. Genomically integrated RAM was quantifiable as it contained a different selectable marker (TmpR).

### RAM can be deployed in a microbial community to capture novel phage hosts

To demonstrate the use of RAM to identify phage hosts in a community context, we infected an 8-member synthetic microbial community with the engineered phage and phagemids (Fig. 2a). In addition to the four microorganisms tested as isolates, we included *Enterobacter cloacae, Salmonella enterica*, and *Shigella sonnei*. No prior studies, to our knowledge, have demonstrated the ability of P1 to infect these three microorganisms. We also included *Pseudomonas aeruginosa* PAO1 as a previous paper reported successful P1 phagemid infection.^25^ The 8-member community captures the majority of gram-negative ESKAPE pathogens, a group of microorganisms responsible for the bulk of multi-drug resistant infections and which may therefore be good targets for phage-based interventions.^30,31^ To discern between these and other microorganisms, RAM provides ∼400 bp sequences that cover the V6-V8 variable regions of the 16S rRNA (Fig. 2b). The microorganisms were mixed at equal OD and then exposed to phage particles for 2 hours.

**Figure 2.**
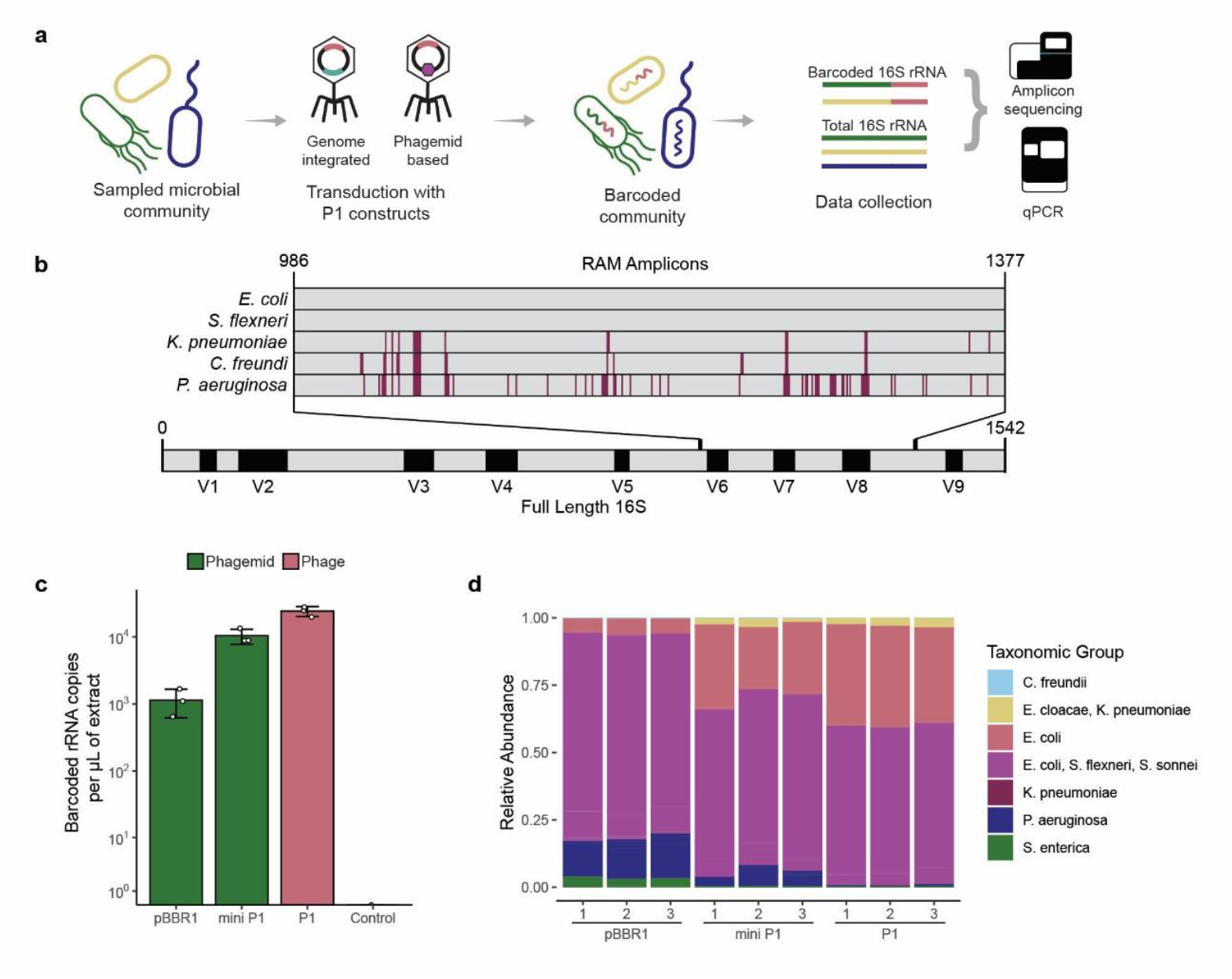
Transduction of a synthetic community by RAM-containing phage particles. **a**, Depiction of the workflow in a microbial community. Biomass from the eight-member community was exposed to phage particles containing RAM constructs for two hours. Total RNA was extracted and analyzed with RT-qPCR for quantification and amplicon sequencing for characterization of both total community and barcoded taxa. **b**, Alignment of Sanger-sequenced, barcoded 16S rRNA from individual species and the location of RAM amplicons within the full length 16S gene. In the amplicons, grey represents nucleotides that match to *E. coli* and red represents nucleotides that differ from *E. coli*. In the full length 16S rRNA gene, black represents the location of highly variable regions within the 16S rRNA gene. **c**, Quantification by RT-qPCR of barcoded 16S rRNA produced through transduction of the synthetic community with phage particles containing the given construct. The control scenario entailed addition of wildtype P1 phage particles that had no incorporation of RAM. Points indicate individual replicates, bars indicate the average value, and error bars the standard deviation. **d**, Relative abundance of different species barcoded by RAM constructs. Reads were assigned using a custom BLAST database generated from publicly available genomes of the different taxa included in the synthetic community. All data was produced with three biological replicates.

Both engineered P1 and phagemids resulted in barcoded 16S rRNA signal when deployed in the synthetic communities, as revealed by RT-qPCR (Fig. 2c). Results showed the most barcoded 16S rRNA for the P1 genomically integrated construct, followed by the mini-P1 phagemid, and then the pBBR1 phagemid (Fig. 2c) (Welch’s, one-tailed, unpaired t-test. *P* = 0.006, 0.012 for P1 > mini P1 and mini P1 > pBBR1, respectively). RAM signal in a community is a combination of barcoded 16S rRNA from the mixture of different phage hosts. Some elements of that signal are species specific, such as the strength of the promoter driving RAM expression.

We then performed amplicon sequencing of the synthetic community’s barcoded 16S to characterize the hosts of the different constructs. Relative abundances of the various species showed consistency between replicates for a single construct but distinct host dynamics across constructs (Fig. 2d). The most abundant group of host species across all constructs consisted of *E. coli, S. flexneri*, and *S. sonnei*. These three organisms were grouped in such a way because the region of the 16S gene adjacent to the RAM barcode is identical in sequence between them (Fig. 2b). A taxonomic group just for *E. coli* was created because a single copy of the 16S gene in *E. coli* is unique enough to distinguish from the *Shigella* species. The same logic applied for the *E. cloacae, K. pneumoniae* group and the individual *K. pneumoniae* group. The phage host community of the pBBR1 phagemid was different from the mini-P1 and P1 communities, while the mini-P1 phagemid and P1 had similar host communities, apart from an additional fraction of *P. aeruginosa* in the mini P1 host community. It should be noted that studies have found that *P. aeruginosa* PAO1 can perform natural transformation, albeit to a limited degree in conditions similar to our experiments, which suggests the possibility that it could be up taking extracellular DNA from the lysate or culture.^32^ We observed barcoding of *S. enterica* 16S rRNA with the pBBR1 phagemid, revealing a novel host of the P1-derived construct and demonstrating RAMs ability to assay host-potential of a phage in a microbial community context.

### A novel order of P1 hosts was revealed in a wastewater microbial community

In our next set of experiments, we deployed the different RAM constructs in a wastewater microbial community derived from wastewater influent sampled from the Upper Brays Wastewater Treatment plant (Fig. 3a). We used the microbial community in untreated wastewater because it contains an abundance of enteric bacteria that P1 is known to infect. As with the synthetic community, all phage constructs resulted in RAM signal, as revealed by both RT-qPCR and amplicon sequencing of the barcoded 16S rRNA. The P1 genomically integrated construct again generated the most barcoded rRNA followed by the mini P1 phagemid and then the pBBR1 phagemid (Fig. 3b; Welch’s, one-tailed, unpaired t-test. *P* = 0.0008, 0.002 for P1 > mini P1 and mini P1 > pBBR1, respectively). Across all constructs, 127 unique amplicon sequencing variants (ASVs) were barcoded, 99 of which also appeared in the native community (Fig. 3c). The fraction of the community not barcoded by RAM (95 ASVs) were either incapable of being infected by any of the constructs or not abundant enough to detect successful infection with our technology. A fraction of ASVs were barcoded that did not appear in the native community (28), likely due to the compositional nature of sequencing data and the displacement of signal from rare ASVs by more abundant ASVs in the native microbial community.

**Figure 3.**
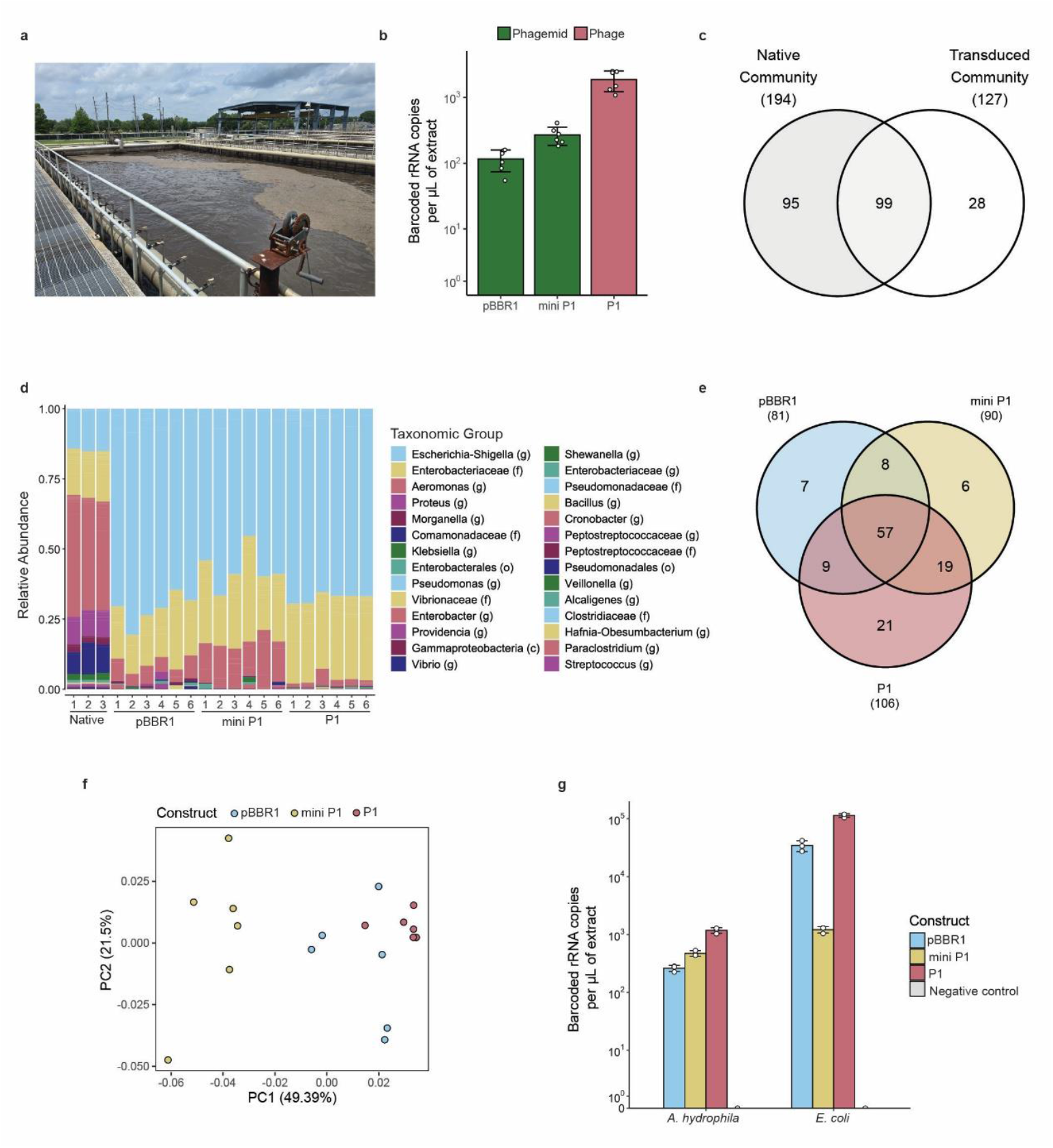
Transduction of a wastewater microbial community by RAM containing phage particles. **a**, Aerobic digester from the Upper Brays Wastewater Treatment plant where wastewater influent was collected. **b**, Quantification by RT-qPCR of barcoded 16S rRNA produced through transduction of the wastewater community with phage particles containing the given construct. Points indicate individual replicates, bars indicate the average value, and error bars the standard deviation for six biological replicates. **c**, Number of unique ASVs barcoded across all of the constructs compared to ASVs in the native microbial community. **d**, Relative abundances of observed taxa in the native community and transduced communities. Letters in parenthesis represent the taxonomic level the group could be defined at, (g) for genus, (f) for family, and (o) for order. **e**, Number of unique, barcoded ASVs shared across different phage and phagemid constructs. **f**, A weighted Unifrac PCoA on communities barcoded by the different constructs. PERMANOVA analysis shows significant clustering of the communities barcoded by the constructs (Fig. S7, q-value < 0.05). **g**, Quantification by RT-qPCR of barcoded 16S rRNA in *A. hydrophila* and *E. coli* MG1655 from different constructs. Points indicate individual

The native community was primarily composed of *Aeromonas* (g), a common genus in wastewater settings, with lesser amounts of *Escherichia-Shigella* (g), *Enterobacteriaceae* (f), *Proteus* (g), and *Comamonadaceae* (f) (Fig. 3d, S5). The phage host ASVs, those ASVs barcoded by RAM in the different constructs, were mainly *Escherichia-Shigella* (g) but also *Enterobacteriaceae* (f) and *Aeromonas* (g) (Fig. 3d). The infection of the *Escherichia-Shigella* (g) and *Enterobacteriaceae* (f) groups aligns with what is known about the host range of P1 and its different constructs. However, *Aeromonas* (g) infection by P1 or its derivatives has not been reported to our knowledge. To confirm our findings, we then screened ten *Aeromonas* species that we isolated from wastewater for infection by P1 constructs. A fraction of the isolates were already resistant to the antibiotics we were using for selective plating (2/10 for trimethoprim, 7/10 for carbenicillin), again highlighting the benefit of using RAM for host identification over selective plating. To confirm via traditional phage-culturing methods, we screened eight of the trimethoprim-susceptible isolates and identified one as also being susceptible to P1 infection (Fig. S6). Whole genome sequencing of the infected host showed the presence of the P1 genome along with the host genome, which we confirmed to be *Aeromonas hydrophila* TN-97-08. Exposure of this isolate to our RAM containing phage particles generated barcoded 16S rRNA for every construct (Fig. 3g). This demonstrates how RAM can be used to identify novel host taxa for phages.

We performed further analysis to assess the degree of similarity of the host communities revealed by RAM for the three constructs. Each construct barcoded similar numbers of ASVs, 81, 90, and 106 for pBBR1, mini P1, and P1, respectively (Fig. 3e). There was a large overlap between all three constructs with 56 ASVs shared between them, and the most shared ASVs between mini P1 and P1, with an additional 19 shared ASVs. A weighted Unifrac analysis on all samples found that replicates from each construct clustered into unique groups (Fig. 3f; PERMANOVA. *q* = 0.005, 0.008, 0.005 for P1/mini P1, P1/pBBR1, and mini P1/pBBR1. See Fig. S7). This suggests that the backbone used to carry RAM significantly influenced the infection efficiency and the host range. Interestingly, pBBR1 clustered more closely to the P1 cluster than the mini P1 cluster, suggesting a higher similarity in host range between these two constructs. The mini P1 host community was distinct from the P1 host community, indicating that a phagemid with the P1 packaging site does not have an identical host range to the P1 phage. (Fig. 3G). These findings show that the phage construct has a significant impact on phage infection efficiency and host range in microbial communities. replicates, bars indicate the average value, and error bars the standard deviation for three biological replicates.

### Two native P1 tail fibers confer a significantly different host range in a wastewater microbial community

To further explore host range determinants in P1, we deployed RAM to characterize the host ranges of two unique tail fibers (S and S’) naturally encoded in an invertible cassette in the P1 genome (Fig. 4a). While the function of every gene in the cassette has not been determined with extreme certainty, it is thought that R attaches tail fibers to the phage base plate, S and S’, are the tail fibers themselves, U and U’, are involved with tail fiber maturation, and cin is the recombinase that inverts the Sv + U and Sv’ + U’ regions.^33^ The tail fibers share an Sc region at the N-terminus, and differ in the inverted Sv or Sv’ regions at the C-terminus. The Sv and Sv’ regions are highly dissimilar (Fig. S8) and are the regions imparting a different host specificity between the S and S’ tail fibers. Previous work suggests sugars in the outer core of the LPS as the target of the S tail fiber and repeated O-antigen units as the target of the S’ tail fiber.^22,34,35^ To apply RAM to assess host range differences imparted by these two different tail fibers, we removed the coding sequences for Sc, Sv, U, Sv’, U’, and cin from the P1 genome.^22^ We then expressed Sc, Sv, U (for S) and Sc, Sv’, U’ (for S’) coding regions from plasmids in separate hosts to generate lysates of phage particles containing only one of the two tail fibers (Fig. 4b).

**Figure 4.**
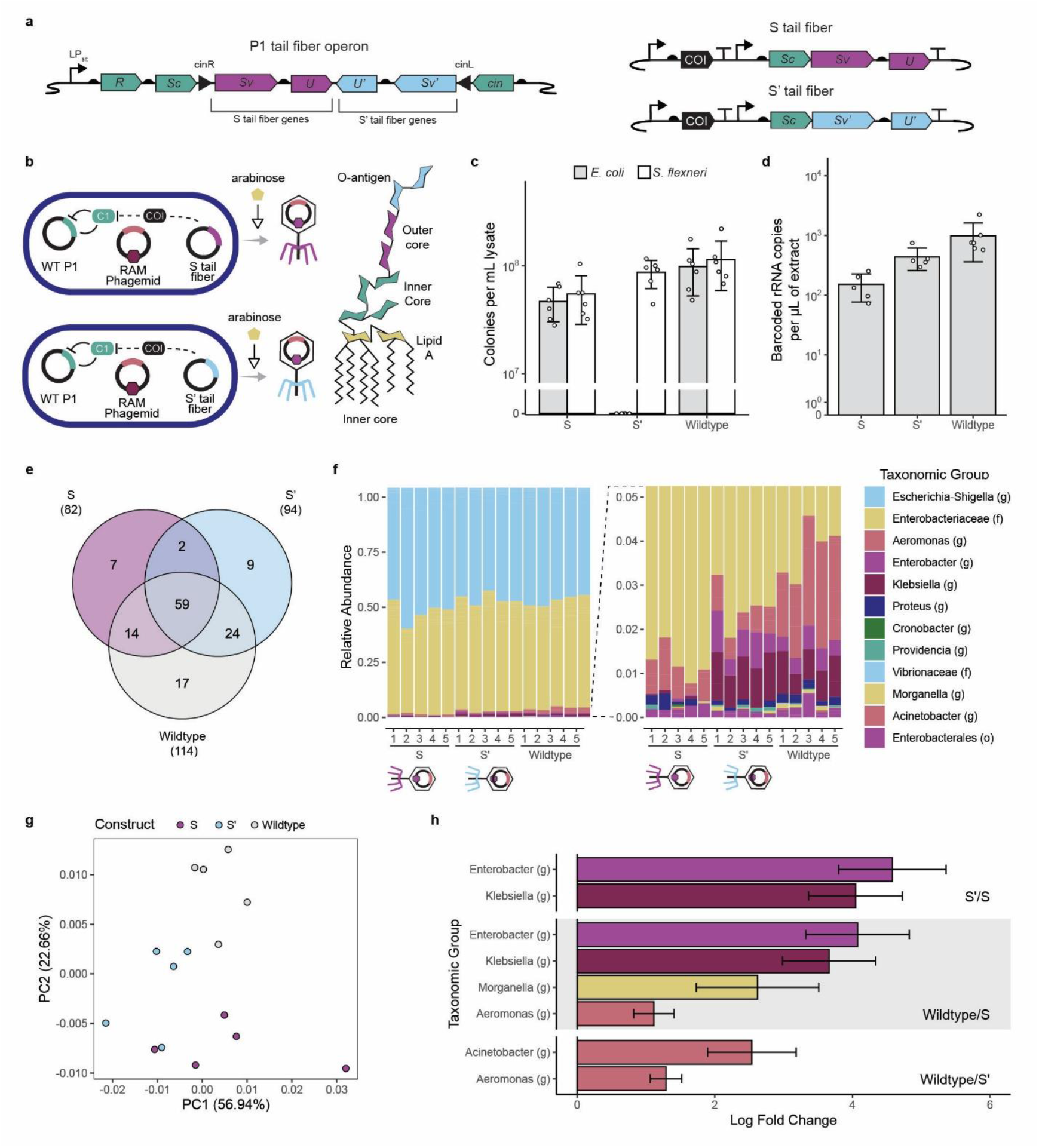
Host range of P1’s two unique tail fibers. **a**, Depiction of the native tail fiber operon in the P1 genome and its reconstitution in plasmids. The R protein attaches the tail fibers to the P1 baseplate. The U and U’ proteins are thought to be tail fiber maturation proteins. Both S and S’ tail fibers contain the Sc region and are differentiated by the unique Sv or Sv’ region that is inverted by the cin gene. The sequence from Sc to cin, including those operons, was removed from the P1 genome. Each unique tail fiber was then reconstituted on different plasmids with either Sc, Sv, U or Sc, Sv’, U’.^22^ **b**, Process for the production of phage particles with unique tail fibers and the tail fibers’ LPS target. The process is the same as for phagemid production, except the P1 lysogen has the tail fiber coding region removed and the lysis plasmid also contains sequences for the appropriate tail fiber. As indicated by the colors, the S tail fiber is though to target sugars in the outer core and the S’ tail fiber the O-antigen. **c**, CFUs generated through selective plating of *E. coli* MG1655 and *S. flexneri* 2a confirms known hosts of P1’s S and S’ tail fibers. Points indicate individual replicates, bars indicate the average value, and error bars the standard deviation for six biological replicates. **d**, Quantification by RT-qPCR of barcoded 16S rRNA produced through transduction of the wastewater community with phage particles containing each tail fiber construct. Wildtype phage particles contain the pBBR1 phagemid generated from a P1 lysogen with a fully intact tail fiber coding region. Points indicate individual replicates, bars indicate the average value, and error bars the standard deviation for five biological replicates. **e**, Number of unique, barcoded ASVs shared across different tail fiber constructs. **f**, Relative abundances of barcoded taxa by the different tail fiber constructs. Letters in parenthesis represent the taxonomic level the group could confidently be defined at, (g) for genus, (f) for family, and (o) for order. On the right is a close up of the relative abundance of the less abundantly barcoded microorganisms. **g**, A weighted Unifrac PCoA on communities barcoded by the different tail fiber constructs. PERMANOVA analysis shows significant clustering of the communities barcoded by the tail fber constructs (Fig. S10, q-value < 0.05). **h**, Log fold change of differentially abundant taxa between the three tail fiber constructs as revealed by ANCOM-BC analysis. Positive values indicate enrichment of a taxonomic group in one set of tail fiber constructs relative to another. Specific tail fiber construct comparisons are indicated in the bottom right hand corner of each section. Bars represent point estimates and error bars the standard error for five biological replicates.

We used the pBBR1 phagemids to deliver RAM to potential hosts.

Initial characterization of our lysates in monoculture via selective plating and RT-qPCR to measure RAM signal showed agreement with previous literature. The S tail fiber can infect both *E. coli* MG1655 and *S. flexneri* 2a 24570, while the S’ tail fiber can only infect *S. flexneri* 2a 24570 (Fig. 4c, S9). The lysate generated using the P1 genome with the wildtype, fully intact tail fiber coding region, demonstrated an infection profile similar to the S tail fiber. However, the plating assay would likely not detect a lower fraction of S’ tail fiber if it was present in this scenario. The natural inversion frequency of the tail fibers is not well understood, nor the specific cellular cues that trigger inversion, but the initial infection of the *E. coli* MG1655 host we used to generate the phage particles with the wildtype tail operon could only have occurred via a P1 particle with an S tail fiber.

As in our previous experiment, we exposed a wastewater microbial community to the phage particles. We observed significantly more barcoded 16S rRNA by the S’ tail fiber construct relative to the S tail fiber (Fig. 4d; Welch’s, one-tailed, unpaired t-test. *P* = 0.043, 0.01 for Wildtype > S’ and S’ > S, respectively). All three tail fiber constructs shared many ASVs (59), and each individual tail fiber construct shared more unique ASVs with the wildtype construct than they did with each other (14 for S and wildtype, 24 for S’ and wildtype versus 2 for S and S’) (Fig. 4e). This suggests that the wildtype construct is a mixture of S and S’ tail fibers as the wildtype also barcoded organisms that were unique to only the S or S’ tail fiber. All three tail fiber constructs barcoded mainly *Escherichia-Shigella* (g) and *Enterobacteriaceae* (f) in roughly equivalent amounts (Fig. 4f). Variable composition between the different tail fiber groups was especially apparent amongst ASVs in lower abundance (Fig. 4f). A PERMANOVA on a weighted Unifrac analysis showed there were significant differences in the community structure of the barcoded ASVs between the three tail fiber constructs (Fig. 4g; PERMANOVA. *q* = 0.039, 0.012, 0.012 for S/S’, S/Wildtype, and S’/Wildtype. See Fig. S10). To identify taxonomic groups that were significantly enriched or depleted between the tail fibers’ host communities, we performed an ANCOM-BC analysis (Fig. 4h). The S’ tail fiber was enriched in *Enterobacter* (g) and *Klebsiella* (g) relative to the S tail fiber. The wildtype tail fibers, relative to the S tail fiber, were also enriched with *Enterobacter* (g) and *Klebsiella* (g), suggesting that a fraction of the wildtype tail fibers were also of the S’ type. The wildtype tail fibers were additionally enriched in *Morganella* (g) and *Aeromonas* (g) relative to the S tail fiber, and *Acinetobacter* (g) and *Aeromonas* (g) relative to the S’ tail fiber.

Together our results demonstrate that P1’s tail fibers significantly influence the host range of a given phage particle. Although our wildtype particles were generated from a P1 lysogen initially in the S tail fiber configuration, our results indicate that tail fibers in the S’ configuration were also generated since hosts unique to the S’ tail fiber were barcoded by wildtype phage particles. These results demonstrate that RAM can be applied to study the viral ecology of the different tail fibers.

## Discussion

RAM is a genetic tool that can rapidly accelerate the study of phage host range in microbial communities. By barcoding host 16S rRNA, RAM takes advantage of widely used molecular biology methods, such as RT-qPCR and amplicon sequencing, to generate both quantitative and qualitative data on phage-host dynamics. The 16S rRNA sequence tagged by RAM can identify a unique host, while the barcode can identify a unique experimental condition. With NGS technologies, many hosts and experimental conditions can be analyzed simultaneously. The total amount of barcoded RNA can then give insight into the series of steps related to phage infection that result in the production of barcoded RNA. Through this streamlined process, RAM addresses shortcomings of many other phage-host characterization approaches, such as a requirement for host culturability, low throughput, difficult implementation, and high expense.

To demonstrate RAMs capabilities, as well as provide a blueprint for how it could be implemented in other phages, we deployed RAM in phage P1, a phage of particular interest in many applied and fundamental contexts. We used the CAST system to incorporate RAM into the P1 genome. Many techniques exist and are being developed to engineer phage genomes, some specifically for lytic phages which are more difficult to manipulate than lysogenic phages.^36–38^ CAST has been deployed in lytic phages and also allows for easy programmability by straightforward manipulation of the gRNA.^27,28^ In some situations a phage may not be amenable to genetic modification, we therefore also demonstrated deployment of RAM in phagemids. Due to their small size, phagemids are easier to modify and incorporate into high-throughput workflows than genomic engineering methods. Additionally, phagemids have already been constructed for phages T7, M13, and Lambda, and tools exist to predict elements of phage packaging that could help with phagemid construction in novel phages.^39–43^ The choice of genomic integration or phagemid appeared to affect total RAM signal, largely due to a lower yield of phagemid containing phage particles relative to phage particles containing genomically integrated RAM.

RAM provided quantitative and qualitative information on the host range and infection dynamics of P1. In monoculture, synthetic community, and wastewater community contexts, P1 with genomically integrated RAM produced the highest amount of barcoded signal. The amount of barcoded 16S rRNA produced is a combination of factors, some of which include initial phage particle concentration, efficiency of binding to cell surface, and efficiency with which the phage surmounts host defenses. If proper controls are implemented, RAM could quantify these and other factors important for phage infection. In the synthetic community, RAM identified a new P1 phagemid host in *S. enterica*, demonstrating its ability to assay host potential of many species at once. RAM could be deployed in other controlled communities to assay, for example, the effect of single gene knockouts in a phage and/or its host in high-throughput. Alternatively, culture collections could be screened with RAM-containing phage(s) to identify potential hosts.^44–46^

We also deployed RAM in a wastewater community without knowing the selection of microorganisms present *a priori*. Our results largely recapitulated what is known about P1 and its phagemid host range while also identifying a completely novel order containing P1 hosts in Aeromonadales. The specific *Aeromonas* (g) hosts observed, includes species that are known human and/or animal pathogens, identifying P1 as a potential tool to fight their infection.^47^ Furthermore, RAM was sufficiently sensitive to identify a significantly different host range conferred by P1s two unique tail fibers. It is important to note that the amount of barcoding and the specific barcoded hosts in these contexts is a function of the underlying microbial community, which can vary widely in wastewater even when wastewater is collected from the same wastewater treatment plant. As shown here, RAM enables the study of host range in unculturable microorganisms and in environmental conditions representative of a phage’s natural environment.

In this approach, phage hosts are identified by sequencing barcoded 16S rRNA. Thus, there are limits in the ability of RAM to distinguish between closely related species and tie specific functions of a species to its 16S sequence, as these are limits of 16S amplicon sequencing generally. Potential solutions could include long reading sequencing to attain a larger fragment of 16S rRNA, targeting a different marker gene for barcoding, or assembly of MAGs to connect 16S sequence to specific traits of the host microorganism.^48–50^ Alternatively, one could express an artificial RNA to act as a synthetic host barcode instead of targeting the native 16S rRNA.

We envision RAM can be applied to understand host range determinants in P1, extended to study other phages, and incorporated into high-throughput workflows. In the case of P1, several different traits likely have outsized impacts on its broad host range, including its own methylation system and its darAB anti-restriction system.^33,51,52^ While we demonstrate RAM in lysogenic P1, there is no reason why it could not be implemented in other groups of phages, such as lytic or RNA phages as RAM works at the level of RNA, a molecule required in the life cycle of all known phages. To fully understand the host range of the 10^7^-10^9^ viral species, high throughput methods need to be developed, especially methods that can train or take advantage of powerful learning tools. Many recent studies rely on plaque assays or selective plating to characterize relationships between phage and host in a semi-high throughput manner, but they could massively benefit from incorporation of RAM and its sequencing-based readout into their workflows.^53–57^ RAM can serve as a high-throughput platform for investigating phage-host interactions and generate large data sets that serve to advance foundational knowledge of phage-host ecology.

## Acknowledgements

We would like to thank the original members of the RAM team not involved with this paper directly, but who made this paper possible by bringing RAM into the world, including Dr. Prashant B. Kalvapalle, August Staubus, Dr. Lauren Gambill, Dr. Kiara Reyes Gamas, and Dr. Li Chieh Lu. We thank Dr. Corwin Miller for providing the initial P1 phage and P1 phagemids that contained the elements incorporated into our genetic constructs. We thank Dr. Shyam Bhakta for allowing us to use his modular cloning system, the Bacterial Toolkit (BTK), to build our genetic constructs. This research was supported by National Science Foundation grants 2227526 (to JC, JJS), 2237052 (to LBS) and 2237512 (to JC). Research was sponsored by the Army Research Office and was accomplished under Cooperative Agreement Number W911NF-24-2-0073. ZWL was supported by funding from the Environmental Research and Education Foundation. SKS was supported by funding from the National Science Foundation Graduate Research Fellowship Program. The views and conclusions contained in this document are those of the authors and should not be interpreted as representing the official policies, either expressed or implied, of the Army Research Office or the U.S. Government.

