## Supplementary material for "Cross-order detection of bacteriophage transduction in communities using ribosomal RNA barcoding": Methods

### Strains, Plasmids, and Cloning

A list of strains can be found in Supplementary Table S1. All strains were cultured in Lysogeny Broth (LB) consisting of 10 g/L tryptone, 5 g/L yeast extract, and 10 g/L NaCl with an additional 15 g/L of agar for plates. Phage lysis media (PLM) consisted of LB with an additional 100 mM $MgCl_2$ and 5 mM $CaCl_2$. M9 minimal medium consisted of 12.8 g/L $NaHPO_4*7H_2O$, 3 g/L $KH_2PO_4$, 0.5 g/L NaCl, 0.4% glucose, 2 mM $MgSO_4$, 0.1 mM $CaCl_2$, and 0.2% casamino acids. Antibiotics were added at concentrations of 100 µg/mL carbenicillin, 20 µg/mL chloramphenicol, 50 µg/mL kanamycin, or 10 µg/mL trimethoprim in liquid culture and on plates. A concentration of 40 µg/mL trimethoprim was used for *A. hydrophila*. Inducers were added at concentrations of 13 mM arabinose for inducing lysis or 100 ng/mL anhydrotetracycline for inducing λ-Red genes in BioDesignER.

A list of plasmids, phagemids, and phages can be found in Supplementary Table S2. Plasmids and phagemids were constructed via Golden Gate assembly or Gibson assembly.[1,2]

Modifications to the P1 genome were implemented through the CAST system or BioDesignER, a λ-Red based system. CAST was used to add RAM and BioDesignER was used to knockout the native tail fiber coding region. A CAST plasmid was constructed containing the full complement of CAST components, including gRNA, Cas/Tn7 proteins, and genetic payload. A gRNA was chosen that would insert the genetic payload away from essential genes. The final location of the genetic payload was upstream of the *mod* gene, a component of the modification system, and downstream of the *lxc* gene, a modulator of the master C1 repressor, both non-essential genes.[3] A depiction of the genetic payload, which includes a single operon to express RAM and TmpR, can be found in Supplementary Figure S1a. The CAST plasmid was transformed into a host that contained the P1 genome as well as a plasmid to induce P1 lysis. A colony from the transformation was used to generate a lysate, protocol detailed below, with a mixture of phage particles containing modified and unmodified P1 genomes. The lysate was then subjected to the transduction to plating protocol using trimethoprim to select for colonies infected with a successfully modified version of P1. The standard protocol for λ-Red based modifications in BioDesignER were followed.[4] Homology arms corresponding to the tail fiber coding regions were taken from prior literature.[5] Oligos that included tails for the homology arms and primer regions corresponding to a TmpR gene were used to generate dsDNA for transformation via PCR. The version of P1 with KanR in the genome called CM01 was a gift from a collaborator.

### Lysate Generation

*E. coli* MG1655 was used as the host strain to generate all phage and phagemid containing particles. The lysates containing the two different tail fibers were generated from the BioDesignER strain, a derivative of *E. coli* MG1655.[4] For RAM encoded in the P1 genome, the host contained RAM encoded P1 and an inducible *coi* plasmid. For RAM encoded on a P1 phagemid, the host contained P1 with a kanamycin resistance marker, the phagemid, and an inducible *coi* plasmid. For tail fiber related experiments, the host contained P1 with the tail fiber coding region knocked out, the phagemid, and an inducible *coi* plasmid that also contained coding regions for either of the two tail fibers. The host was cultured overnight directly from a glycerol stock or from a colony with appropriate antibiotics. Saturated cultures were then subcultured 100x into 2 mL of LB media containing appropriate antibiotics. After 2 hours, subcultures were pelleted by centrifugation at 4100xg for 10 minutes. Pellets were resuspended without antibiotics in 2 mL of PLM. Arabinose was added at a final concentration of 13 µM to induce the lytic cycle. Cultures were given 2 hours to fully lyse and then mixed with chloroform at a final concentration of 2.5%. Lysates were then vortexed for 30 seconds and centrifuged for 2 minutes at 12,200xg. Supernatant was removed and stored at 4 °C. All growth and lysis steps occurred in an incubator kept at 37 °C and 250 RPM.

We included a PEG concentration step for the lysates containing the S and S' tail fibers as the modifications appeared to lower the number of functional phagemid particles. Cultures were scaled up to 50 mL following our previous lysate preparation protocol. After the 2 hour lysis step, 2.5% chloroform and 0.3 M NaCl (including NaCl in PLM) were added directly to 50 mL flasks containing lysate and allowed to shake at 37 °C and 250 rpm for 15 minutes. Lysates were then centrifuged at 4100g for 30 minutes, and the supernatant carefully collected. PEG 6000 was then added to the lysates to a final concentration of 4g/100mL and the lysates allowed to sit overnight at 4 °C. The lysates were then spun down at 4100g for 45 minutes at 4 °C. The supernatant was poured off and 500 µL of SM buffer added to the pellet. The pellet was resuspended by gentle, up and down pipetting and then allowed to sit for 2 hours. The resuspension was then filtered through a 0.22 µM filter and stored at 4 °C until use.

### Transduction for Plating

Host organisms were cultured overnight to stationary phase directly from a glycerol stock. Hosts were then resuspended to OD 0.5 in PLM and placed on ice. Meanwhile, a dilution series of the phage/phagemid lysates was prepared with PLM in a 96 well plate. Next, 50 µL of host suspension was mixed with 50 µL of phage/phagemid dilution in a different 96 well plate. The plate was then incubated at 37 °C and 250 RPM for 1 hour. Finally, 5 µL of host and phage/phagemid mixture was plated on antibiotic plates to select for appropriate subpopulations of phage/phagemid. Colonies were then counted at the lowest dilution with visibly distinct colonies.

### Transduction for RT-qPCR

Similar to transduction efficiency, host organisms were cultured overnight to stationary phase directly from a glycerol stock. Hosts were then resuspended to OD 0.5 in PLM for all wastewater and Aeromonas experiments and M9 supplemented with 1% casamino acids and 0.4% glucose for the initial monoculture and synthetic community experiments. To create the synthetic community, equal volumes of host suspension were combined together. Phage lysates were spiked at a ratio of 1:10 of host suspension. The mixture was then incubated at 37 °C and 250 RPM for 2 hours before harvesting for RNA extraction.

### RNA or DNA Extraction

RNA was extracted on a Promega Maxwell RSC 48 with a modified protocol for the Maxwell RSC Purefood GMO and Authentication Kit. Modifications are centered on the lysis step. Biomass equivalent to 1 mL of 0.4 OD culture was pelleted and resuspended in 1 mL of CTAB buffer. Samples were then incubated for 5 minutes at 90 °C. Next, samples were removed and given roughly 5 minutes to cool at room temperature before adding 40 µL of Proteinase K. Finally, samples were incubated for 10 minutes at 70 °C before being added to Maxwell cartridges and continuing with the standard kit pipeline. Samples were eluted into 50 µL of elution buffer and immediately processed to remove DNA.

DNA from *A. hydrophila* containing P1 was extracted on a Promega Maxwell RSC 48 with the standard DNA extraction protocol from the RSC Purefood GMO and Authentication Kit. Biomass equivalent to 1 mL of 0.4 OD culture was used in the extraction.

### DNA Removal

DNA was removed using the Invitrogen TURBO DNA-free Kit with a modified protocol. All reactions consisted of 30 µL sample extract, 13 µL $H_2O$, 5 µL 10X TURBO DNase buffer, and 2 µL TURBO DNase. Reactions were incubated at 37 °C for 1 hour after which 5 µL of DNase Inactivation Reagent was added. Reactions were then incubated for an additional 10 minutes at 37 °C with brief inversions/flicking of tubes every 2.5 minutes to resuspend settled DNase Inactivation Reagent. Samples were stored at -80 °C until further processing for RT-qPCR or RT-PCR.

### Reverse transcription quantitative PCR (RT-qPCR)

Primers and probes for RT-qPCR can be found in Supplementary Table S3. Reactions were prepared with PCR Biosystem's qPCRBIO Probe 1-Step Go reaction mix and run on a Quantstudio-3 qPCR thermocycler. A single 10 µL reaction consisted of 0.4 µL of $H_2O$, 5 µL of probe master mix, 0.1 µL of reverse transcriptase, 0.5 µl of a previously prepared primer/probe stock solution, and 4 µL of RNA extract. The final concentration of oligos in the reaction was 0.4 µM for primers and 0.2 µM for probes. RNA extracts were diluted 10x prior to addition to the reaction to minimize the effect of inhibitors. RT-qPCR data from the thermocycler was first analyzed in Thermofisher's Design and Analysis Software 2.7.0. Standard curves were generated from a dilution series of barcoded RAM amplicons from *E. coli* MG1655 that had been converted to DNA through RT-PCR. Concentration of the undiluted DNA standard was determined via qubit. New standard curves were generated for each preparation of primer/probe stock solution. The thresholds determined in the standard curve for a primer/probe stock preparation were manually set to the same value in all RT-qPCR runs using that stock preparation.  All RT-qPCR data in a specific figure panel was generated on the same RT-qPCR run.

### Reverse transcription PCR (RT-PCR)

Primers for RT-PCR can be found in Supplementary Table S3. Reactions were prepared with New England Biolab's LunaScript® Multiplex One-Step RT-PCR Kit on either a BioRad C1000

Touch or T100 thermocycler. A single 25 µL reaction consisted of 8.75 µL $H_2O$, 1.25 µL DMSO, 5 µL of reaction buffer, 1 µL of enzyme mix, 5 µL of a previously prepared primer stock solution, and 4 µL of RNA extract. The final concentration of primers in the reaction was 0.5 µM. RNA extracts were diluted 10x prior to addition to the reaction to minimize the effect of inhibitors. Reactions were scaled up to 50 µL for RAM amplicons to increase the amount of DNA generated. Reactions were subjected to gel electrophoresis and appropriately sized bands extracted with New England Biolab's Monarch® Spin DNA Gel Extraction Kit. Extracted DNA was then sent to Genewiz for sequencing through their Amplicon-EZ pipeline.

### Amplicon sequencing analysis

A depiction of the workflow for amplicon sequencing analysis can be found in Supplementary Figure S11. Sequencing analysis was conducted in Qiime 2 2024.10.[6] Data for barcoded 16S and whole community 16S were initially imported into Qiime separately due to their unique trimming requirements. The Cutadapt plugin was used to trim the forward primer (5'-GCAACGCGAAGAACCTTACC) for both data sets and the unique reverse primer for each data set, (5'-AAGTCATGCCGTTTCATGTGATC) for barcoded 16S and (5'-TGACGGGCGGTGWGTRCA) for the whole community 16S.[7] A residual barcode sequence was trimmed from the barcoded 16S data set (5'-CGGATAACGGGAAAAGCATTGAACACCAT). To make the sequence comparisons between the two datasets equivalent, an additional sequence was trimmed from the whole community 16S data set (5'-AGGCCCGGGAACGT). Primers and the extra sequence from the whole community 16S data set were required to have at least 5 nt aligning to the end of the read and 10 nt for the residual barcode sequence. Primers and the residual barcode sequence were also required to have less than 10% sequence mismatch and 14% sequence mismatch for the extra sequence from the whole community 16S data set. Reads that did not contain these sequences were discarded. The two data sets were then exported from Qiime, combined, and imported back into Qiime together. The DADA2 plugin was used to denoise, merge paired-end sequences, dereplicate, and then filter chimeras.[8] ASVs generated from this process that did not occur more than three times across two samples were removed, along with any ASVs below 250 bp. Taxonomy was assigned via the *feature-classifier sklearn-classify* command using the Silva 138 99% OTU database.[9] Taxonomy for the synthetic community was assigned with the *feature-classifier blast* command using a custom BLAST database constructed from the genomes of the included organisms.

### Statistical methods

Statistical analysis of sequencing data was conducted in Qiime 2 2024.10 using default settings unless otherwise specified.[6] Diversity metrics were calculated with the *diversity core-metrics-phylogenetic* command. The resulting PCoA of the weighted Unifrac analysis was exported from Qiime 2 and visualized in R.[10] The ANCOM-BC analysis was conducted with the *composition ancombc* command.[11] The resulting Qiime 2 files were exported and visualized in R. ANCOM-BC results were only reported in the main text if they were statistically significant (q-value < 0.05).
