## Supplementary Figures for "Cross-order detection of bacteriophage transduction in communities using ribosomal RNA barcoding"

| Item | Description | Page |
|---|---|---|
| Fig. S1 | Architecture of additional genetic constructs | 1 |
| Fig. S2 | Characterization of the different types of phage particles in the lysate | 2 |
| Fig. S3 | Effect of incubation time on the RAM signal | 3 |
| Fig. S4 | Effect of lysate concentration on RAM signal | 4 |
| Fig. S5 | Composition of native wastewater community | 5 |
| Fig. S6 | Selective plating of *A. hydrophila* TN-97-08 | 6 |
| Fig. S7 | PERMANOVA results on the PCoA of the weighted Unifrac distance matrix between different phagemid and phage constructs | 7 |
| Fig. S8 | Amino acid alignment of P1's two unique tail fibers | 8 |
| Fig. S9 | Barcoded 16S rRNA by different tail fiber constructs in monoculture | 9 |
| Fig. S10 | PERMANOVA results on the PCoA of the weighted Unifrac distance matrix between different tail fiber constructs | 10 |
| Fig. S11 | Workflow for processing sequencing data | 11 |

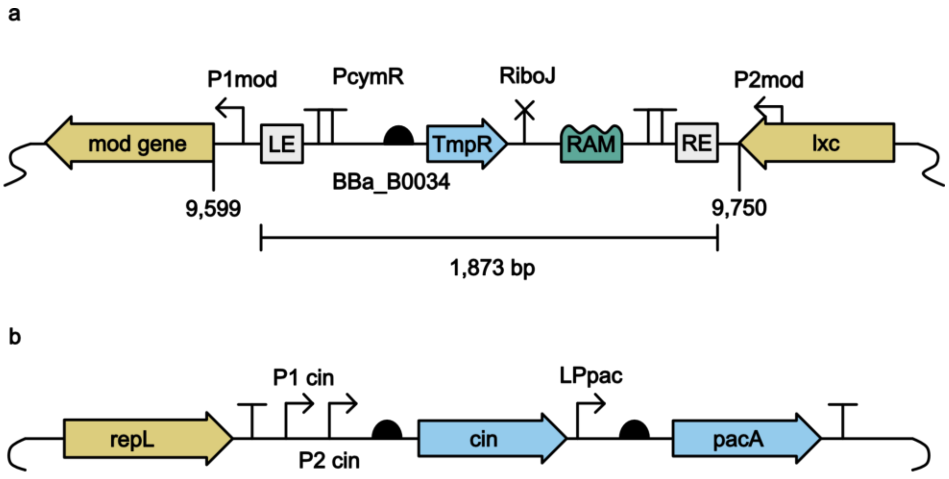


**Supplementary Figure S1. Architecture of RAM insert in the P1 genome and the P1 packaging cassette included in P1 phagemids. a,** Location and architecture of the RAM cassette added to the P1 genome. The *mod* and *lxc* genes are considered non-essential.[1] LE and RE correspond to the left and right transposon ends that are components of the CAST system. Insulators were included on either side of the RAM operon to prevent unwanted inward or outward transcription. Trimethoprim resistance was chosen because it has a relatively small genetic footprint. RiboJ was included to cleave the transcript to prevent unwanted translation of RAM or interference of RAM splicing by additional RNA. **b,** Architecture of the packaging site incorporated into phagemids. Native expression elements were included where needed. The *repL* sequence contains no native or synthetic expression elements, but does still contain the lytic origin of replication. The *repL* protein is expressed from the P1 lysogen. There is no known role of *cin* in phage packaging, but it's inclusion in phagemids has been shown to improve their packaging.[2] The *pacA* protein is a subunit of the pacase enzyme that packages DNA into the phage head.[3] The *pacA* coding sequence also contains the sequence for the packaging site recognized by packaging machinery.

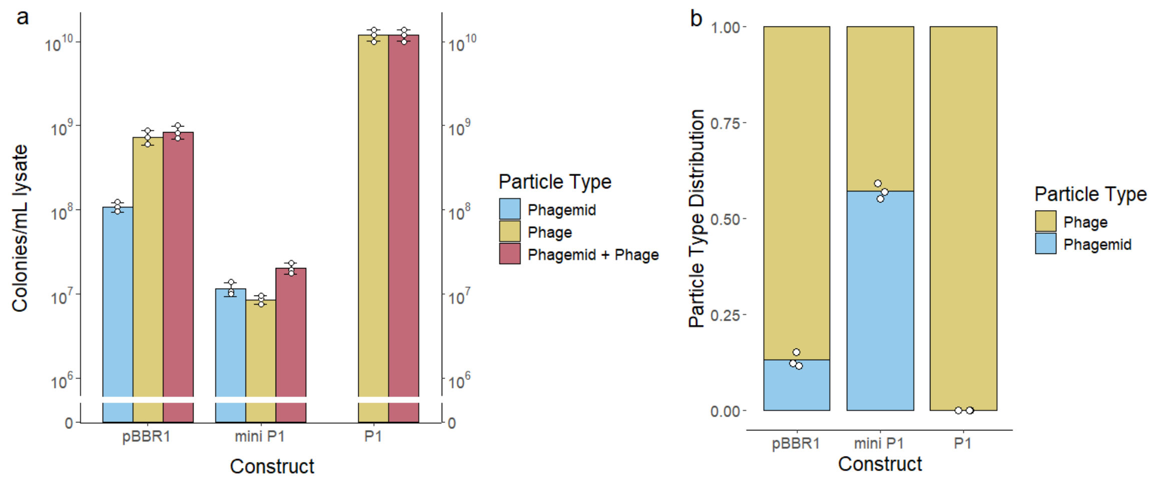


**Supplementary Figure S2. Distribution and characterization of phage particle types in the phagemid lysates. a,** Total CFUs generated by different particle types in the lysate. Different particle types were enumerated by transduction to selective plating with appropriate antibiotics using an *E. coli* MG1655 host. For the phagemids, a version of P1 without RAM but containing an extra KanR gene was used to allow for selection of near-wildtype P1 particles. The condition described as "P1" depicts a lysate generated from a host that contained no phagemids but did contains the same KanR version of P1. The phagemid lysates are mixtures of phagemid and P1 genome containing phage particles. The phagemid production process reduced the total amount of phage particles. **b,** The distribution of different particle types contained in the different lysates. The data shown here was generated with data from Fig. S2a. For both panels, points indicate individual replicates, bars indicate the average value, and error bars the standard deviation (of panel a) of three biological replicates.

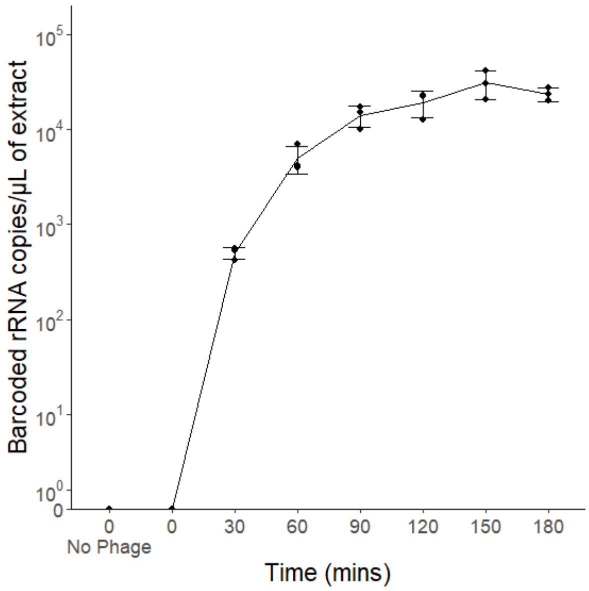


**Supplementary Figure S3. Effect of incubation time on the RAM signal.** P1 phage with genomically encoded RAM was added to a suspension of *E. coli* MG1655 at time zero, and barcoded rRNA was quantified using RT-qPCR in samples of the suspension collected every 30 minutes. As a control, RAM signal was also measured from *E. coli* without the addition of phages (no signal was detected). Based on this data, an incubation time of two hours was selected to incubate RAM containing phages in bacterial cultures before harvesting RNA.

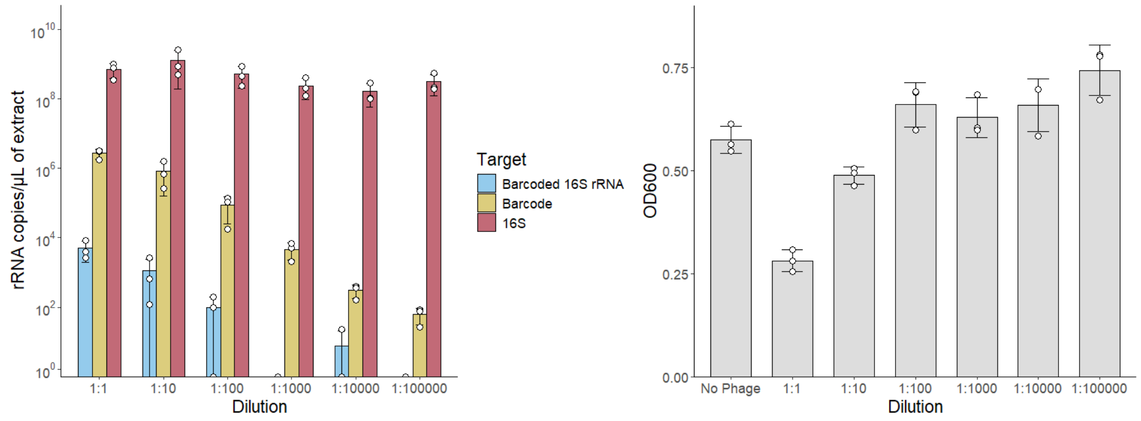


**Supplementary Figure S4. Effect of lysate concentration on RAM signal. a,** Phage lysate of P1 with genomically encoded RAM was exposed to *E. coli* MG1655 at different dilutions to determine the appropriate concentration to add to cultures. The x-axis indicates a particular dilution while the y-axis indicates the total copies of that RNA molecule as measured by RT-qPCR. Different colored bars indicate different targets. The "Barcoded 16S rRNA", is the spliced product and target reported in all other the RT-qPCR based figures. The "Barcode" target indicates the total concentration of barcode including the barcode in spliced products and the unspliced ribozyme. The "16S" target indicates the total concentration of 16S rRNA, including 16S rRNA in the spliced product. 16S rRNA in the spliced product will be a negligible amount relative to the amount of unspliced 16S rRNA. Based on this data, undiluted lysates (1:1), were used in further experiments as dilution only decreased the signal from barcoded 16S rRNA. The concentration of phage particles in both phagemid lysates was less than for P1 with genomically encoded RAM (Fig. S2a) and we therefore assumed no dilution would be the optimal conditions for phagemid lysates as well. **b,** OD600 of the *E. coli* MG1655 cultures at the time of harvesting for RT-qPCR analysis in Fig. S4a. The 1:1 and 1:10 dilutions appeared to inhibit bacterial growth, potentially due to lysis from phage particles.

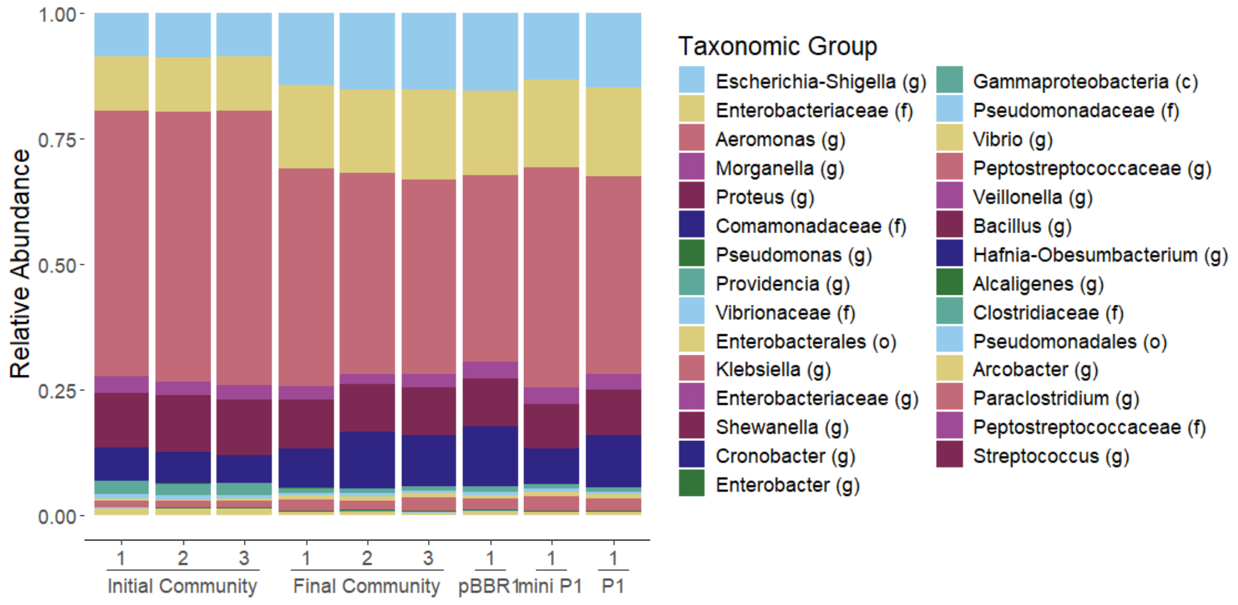


**Supplementary Figure S5. Composition of the native wastewater microbial community.** The initial community is the composition of the wastewater community before exposure to phage particles. The final community is the composition of the wastewater community exposed to wildtype P1 particles. The other columns are the compositions of the wastewater communities exposed to the different constructs (pBBR1 phagemids, mini P1 phagemids, RAM encoded P1). Composition differences between the initial community and the other communities could be due to the addition of phages or growth of the microorganisms during the two hour exposure period. There appeared to be no differences between the wastewater communities exposed to the different types of phage particles, though the limited number of replicates prevents a statistical analysis.

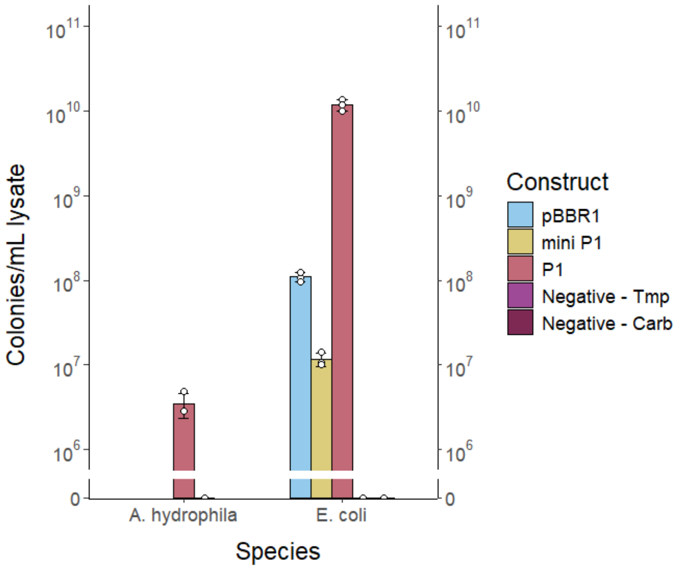


**Supplementary Figure S6. Selective plating of *A. hydrophila* TN-97-08.** We exposed *A. hydrophila* to RAM encoded P1 and plated it  using trimethoprim selection to determine it was susceptible to P1. Selective plating could not be used to determine the susceptibility of *A. hydrophila* to P1 phagemids because the isolate was already resistant to carbapenicillin, the antibiotic we were using to select for phagemids.

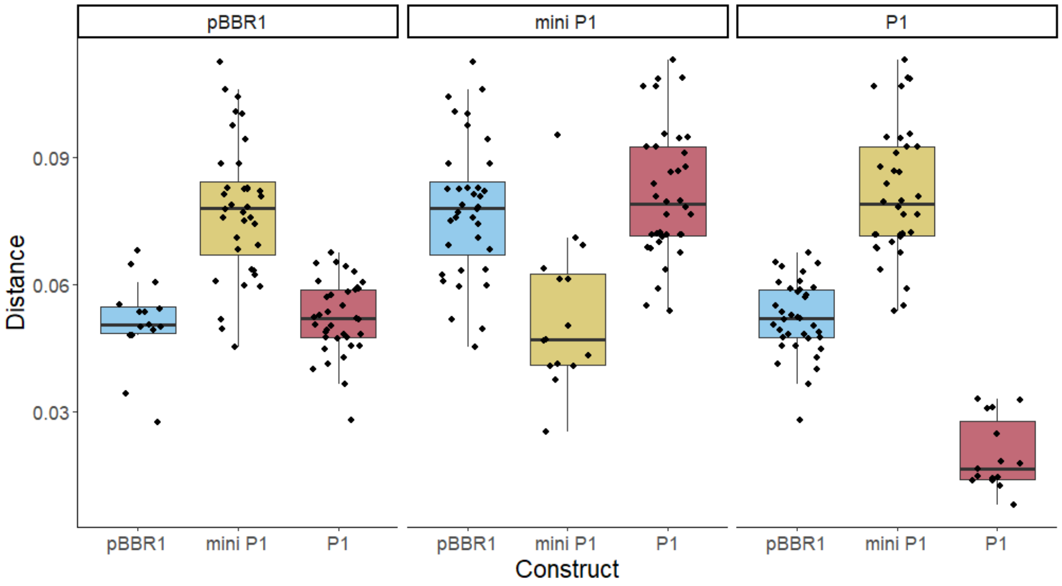


| Group 1 | Group 2 | Sample size | Permutations | pseudo-F | p-value | q-value |
|---|---|---|---|---|---|---|
| **P1** | **miniP1** | 12 | 999 | 18.198388 | 0.003 | 0.0045 |
| | **pBBR1** | 12 | 999 | 5.553165 | 0.008 | 0.0080 |
| **miniP1** | **pBBR1** | 12 | 999 | 8.063046 | 0.002 | 0.0045 |

**Supplementary Figure S7. Distinct host communities of P1 constructs were observed based on PERMANOVA on the PCoA of the weighted Unifrac distances.** Each point represents the distance between two points on the PCoA. Each panel represents distances of the points from the construct named in the panel to points in the groups indicated on the bottom x-axis. The panels contain the distance between the points within that construct (i.e. the first box plot on the left contains the distances between all combinations of pBBR1 points) as well as the distance between the points from the construct named in the panel to the one named on the bottom x-axis (i.e. the second box plot on the left contains the distances between every combinations of two points from pBBR1 and mini P1). Boxplots include the median (central grey line), 25th quartile (bottom of the box), 75th quartile (top of the box), lower whisker (bottom vertical line), and upper whisker (upper vertical line). Hinges extend to the furthest data point out that is no more than 1.5*IQR (Interquartile distance) from the closest quartile. The pseudo-F is a comparison of the distances between points from one group, to distances between points from the two groups combined and is therefore, a measure of the effect size. The q-value is an adjusted p-value that accounts for multiple comparisons and is also a measure of statistical significance.

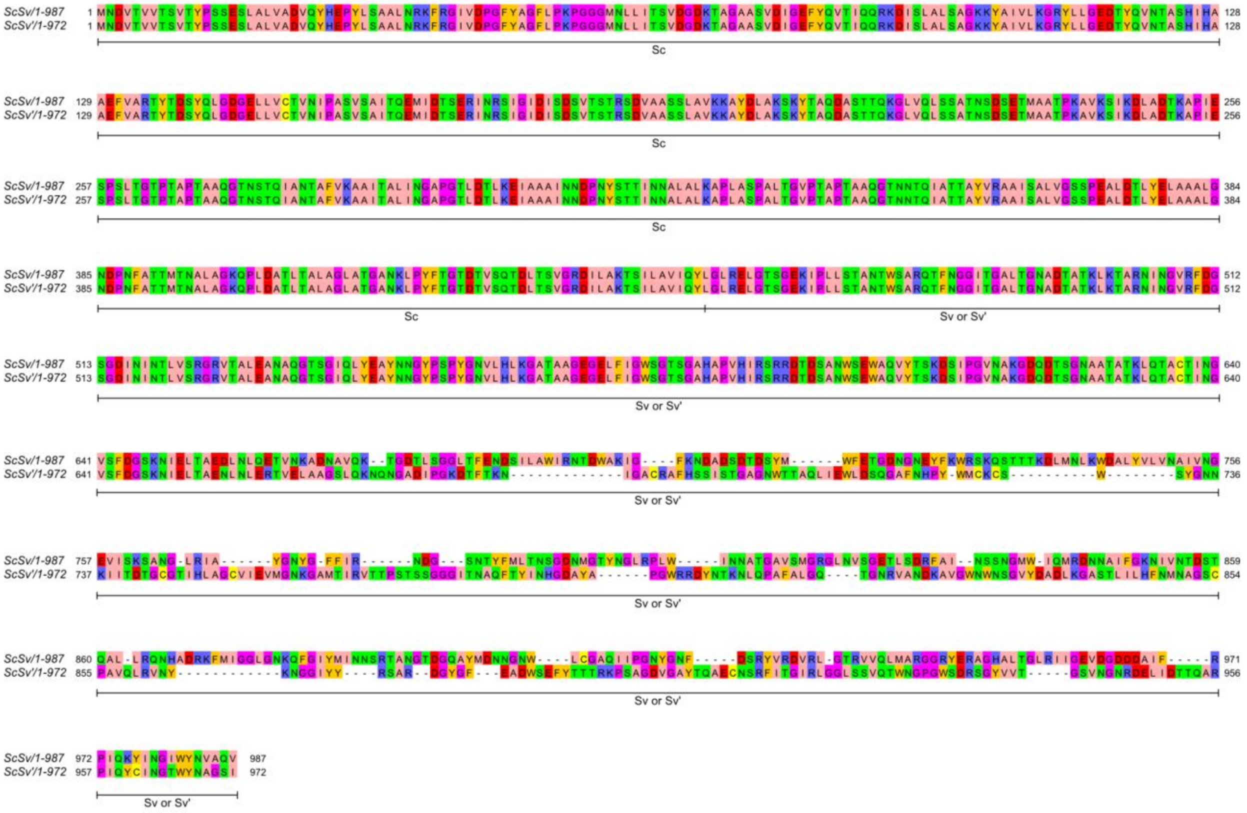

**Supplementary Figure S8. Amino acid alignment of P1's two unique tail fibers.** Both S and S' tail fibers share a tail fiber region called Sc that is not involved in the invertible cassette. The invertible cassette flips either Sv (for S) or Sv' (for S') into position. Both Sv and Sv' share amino acid composition at the N-terminus but have different amino acid profile at the C-terminus. Full S or S' tail fibers are trimers with the Sv and Sv' regions determining interactions with the host LPS.[4]

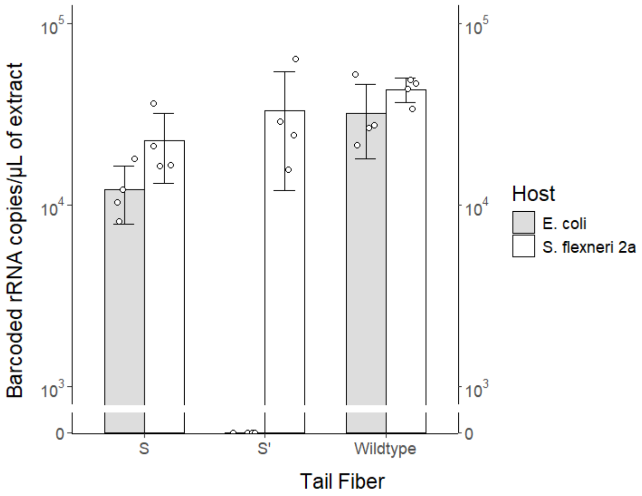


**Supplementary Figure S9. Barcoded 16S rRNA by different tail fiber constructs in monoculture confirms known host specificity of S and S' tail fibers.** A transduction to RT-qPCR assay was carried out using the tail fiber constructs to infect *E. coli* and *S. flexneri* 2a. Results agree with the literature and selective plating data that S can infect *E. coli* and *S. flexneri* 2a, while S' can only infect *S. flexneri* 2a.

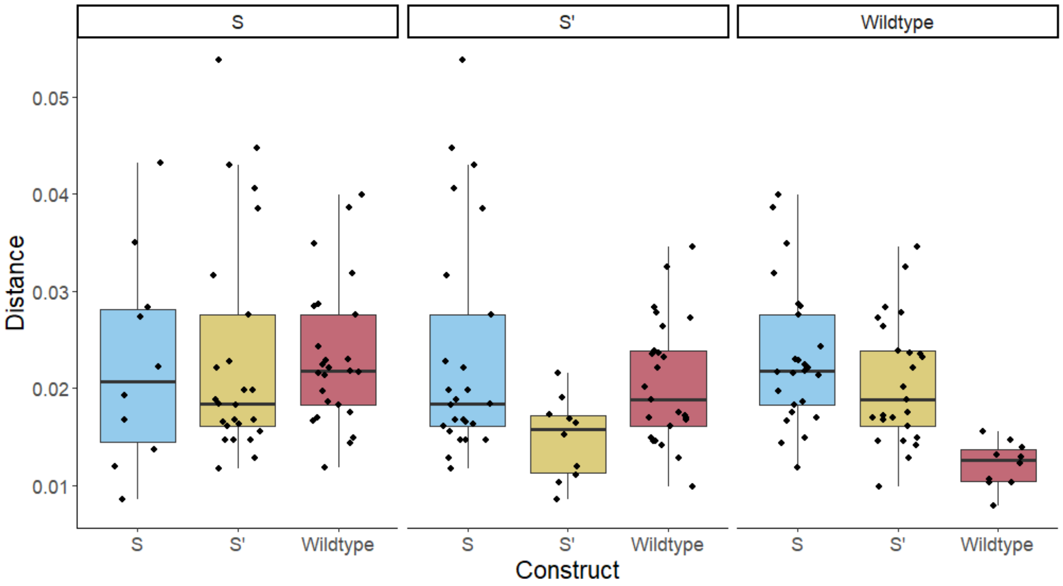


| Group 1 | Group 2 | Sample size | Permutations | pseudo-F | p-value | q-value |
|---|---|---|---|---|---|---|
| **S** | **S’** | 10 | 999 | 3.958099 | 0.039 | 0.039 |
| | **Wildtype** | 10 | 999 | 3.620586 | 0.008 | 0.012 |
| **S’** | **Wildtype** | 10 | 999 | 7.888064 | 0.005 | 0.012 |

**Supplementary Figure S10. PERMANOVA on the PCoA of the weighted Unifrac distance matrix showed statistically significant differences in community composition between each tail fiber construct.** Each point represents the distance between two points on the PCoA. Each panel represents distances of the points from the construct named in the panel to points in the groups indicated on the bottom x-axis. The panels contain the distance between the points within that construct (i.e. the first box plot on the left contains the distances between all combinations of S points) as well as the distance between the points from the construct named in the panel to the one named on the bottom x-axis (i.e. the second box plot on the left contains the distances between every combinations of two points from S and S’). Boxplots include the median (central grey line), 25th quartile (bottom of the box), 75th quartile (top of the box), lower whisker (bottom vertical line), and upper whisker (upper vertical line). Hinges extend to the furthest data point out that is no more than 1.5*IQR (Interquartile distance) from the closest quartile. The pseudo-F is a comparison of the distances between points from one group, to distances between points from the two groups combined and is therefore, a measure of the effect size. The q-value is an adjusted p-value, that accounts for multiple comparisons and is also a measure of statistical significance.

**Supplementary Figure S11. Workflow for processing sequencing data**. The workflow used to analyze amplicon sequencing data. Individual steps are described in grey boxes. Grey boxes include a short description and relevant Qiime 2 commands in italics. Each step will generate files depicted in the black boxes

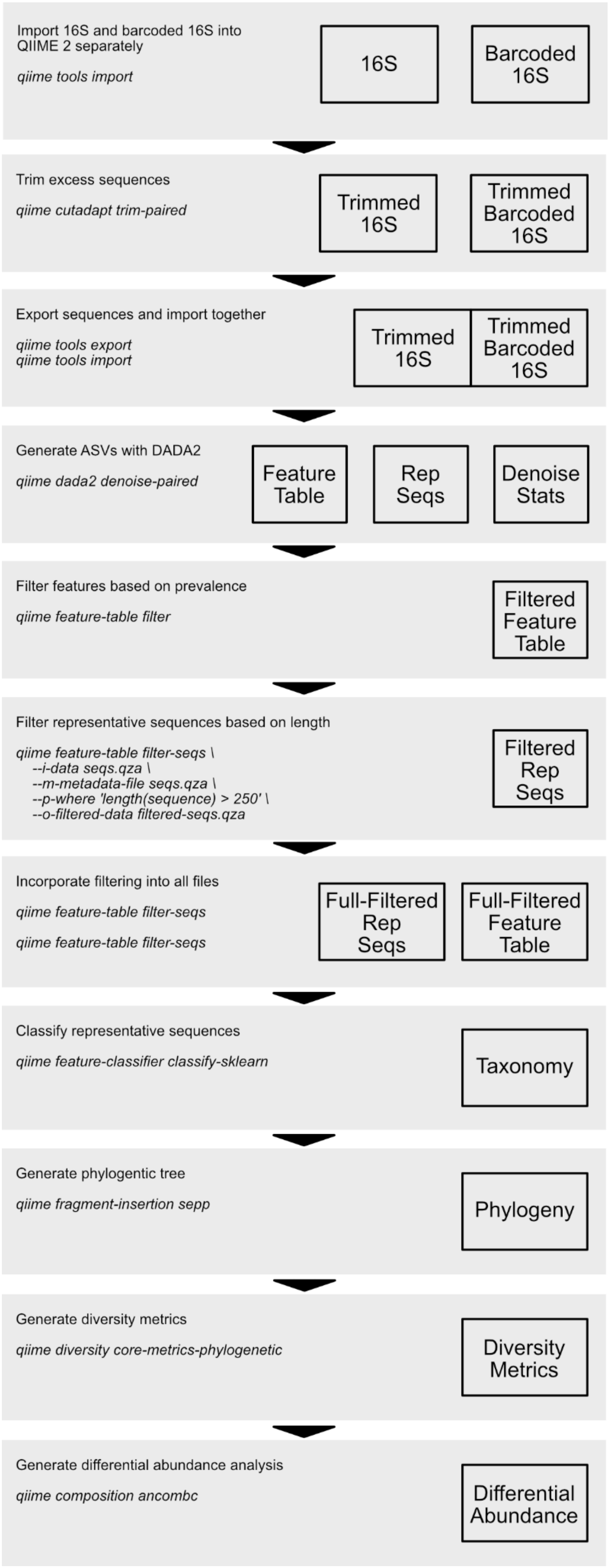
